# Differential impact of reward and punishment on functional connectivity after skill learning

**DOI:** 10.1101/243899

**Authors:** Adam Steel, Edward H. Silson, Charlotte J. Stagg, Chris I. Baker

**Affiliations:** Wellcome Centre for Integrative Neuroimaging, FMRIB, Nuffield Department of Clinical Neurosciences, University of Oxford, Oxford, OX3 9DU, UK; Laboratory of Brain and Cognition, National Institute of Mental Health, National Institutes of Health, Bethesda, MD, 20814; Oxford Centre for Human Brain Activity (OHBA), Wellcome Centre for Integrative Neuroimaging, University Department of Psychiatry, University of Oxford, Oxford, UK, OX3 9DU, UK

**Author notes:** Joint Senior Authors (Charlotte J. Stagg, Chris I. Baker). Corresponding Author: Adam Steel, FMRIB Centre, John Radcliffe Hospital, Headington, Oxford, OX3 9DU, United Kingdom. Competing financial interests: The authors declare no competing financial interests.

**Keywords:** Sequence learning, Motor learning, Consolidation, Motivation, Premotor cortex

## Abstract

Reward and punishment shape behavior, but the mechanisms underlying their effect on skill learning are not well understood. Here, we tested whether the functional connectivity of premotor cortex (PMC), a region known to be critical for learning of sequencing skills, is altered after training by reward or punishment given during training. Resting-state fMRI was collected in two experiments before and after participants trained on either a serial reaction time task (SRTT; n = 36) or force-tracking task (FTT; n = 36) with reward, punishment, or control feedback. In each experiment, training-related change in PMC functional connectivity was compared across feedback groups. In both tasks, reward and punishment differentially affected PMC functional connectivity. On the SRTT, participants trained with reward showed an increase in functional connectivity between PMC and cerebellum as well as PMC and striatum, while participants trained with punishment showed an increase in functional connectivity between PMC and medial temporal lobe connectivity. After training on the FTT, subjects trained with control and reward showed increases in PMC connectivity with parietal and temporal cortices after training, while subjects trained with punishment showed increased PMC connectivity with ventral striatum. While the results from the two experiments overlapped in some areas, including ventral pallidum, temporal lobe, and cerebellum, these regions showed diverging patterns of results across the two tasks for the different feedback conditions. These findings suggest that reward and punishment strongly influence spontaneous brain activity after training, and that the regions implicated depend on the task learned.

---

The potential to use reward and punishment, collectively referred to as valenced feedback, during training has been pursued in recent years as a potential method to increase skill learning and retention (Abe et al., 2011; Galea et al., 2015; Steel et al., 2016; Wachter et al., 2009). Prior behavioral studies of motor adaptation suggest that reward and punishment have differing effects on motor learning. For example, punishment increased learning rate in a cerebellar-dependent motor adaptation task (Galea et al., 2015), while reward prevented forgetting after adaptation (Galea et al., 2015; Shmuelof et al., 2012). Reward may also restore adaptation learning in patients with cerebellar degeneration (Therrien et al., 2016) and stroke (Quattrocchi et al., 2017). Beyond adaptation tasks, in other skill-learning contexts it has been reported that reward improves memory retention compared to punishment (Abe et al., 2011), though these results are somewhat inconsistent across the literature (Steel et al., 2016).

One explanation for the differential effects of reward and punishment on behavior is the recruitment of core set of brain regions involved in feedback processing. It has been suggested that punishment leads to the recruitment of fast learning systems [e.g. medial temporal lobe (MTL)], while reward recruits slow learning systems [e.g. caudate via dopaminergic signaling (Peterson and Seger, 2013b; Wachter et al., 2009)]. In support of this hypothesis, functional imaging studies where fMRI data was acquired concurrent with task performance have reported that reward increases caudate activity in a behaviorally-relevant manner (Peterson and Seger, 2013b; Wachter et al., 2009). In contrast, punishment increases activity in the anterior insula (Shigemune et al., 2014; Wachter et al., 2009) and MTL (Murty et al., 2012b, 2016). However, these studies were primarily conducted during tasks that are statistical in nature. It is therefore not clear whether reward and punishment engage different brain regions depending on the demand of the task being performed

To address this issue, in this study we examined whether the brain regions affected by training with reward or punishment are common across tasks by examining changes functional connectivity induced by training. Because the ‘state’ of the participant is consistent in the pre- and post-training resting-state scans, this technique also allows us to assess the effect of training across two tasks that have very different low-level demands. By comparing the impact of feedback on the change in resting-state functional connectivity after training between the two tasks, we can isolate any task-general effects of feedback without the confound of task performance (i.e. movement). Notably, changes in resting-state functional connectivity after training likely reflect offline-memory processing (Sami and Miall, 2013; Sami et al., 2014) as well as latent effects of task performance including rumination and homeostatic plasticity (Gregory et al., 2016). Therefore, while we cannot attribute any changes in resting-state functional connectivity to offline-processes related to memory, per se, with this approach we can detect the overall impact of feedback on the brain after training.

In the first experiment, based on the well described role of fast and slow learning systems in the context of the serial reaction time task [SRTT; (Doyon et al., 2018)], we examined the impact of feedback valence on neural activity induced by SRTT training. In the second experiment, to test whether any regions impacted by training with feedback generalize to a different motor sequencing task with distinct task demands, we implemented the force tracking task (FTT) with reward and punishment. In both experiments, before and after training we collected 20-minutes of resting-state fMRI data (Figure 1a-d). We have previously presented our behavioral results, which suggested that feedback differentially impacts performance during learning in these two tasks (Steel et al., 2016).

**Figure 1.**
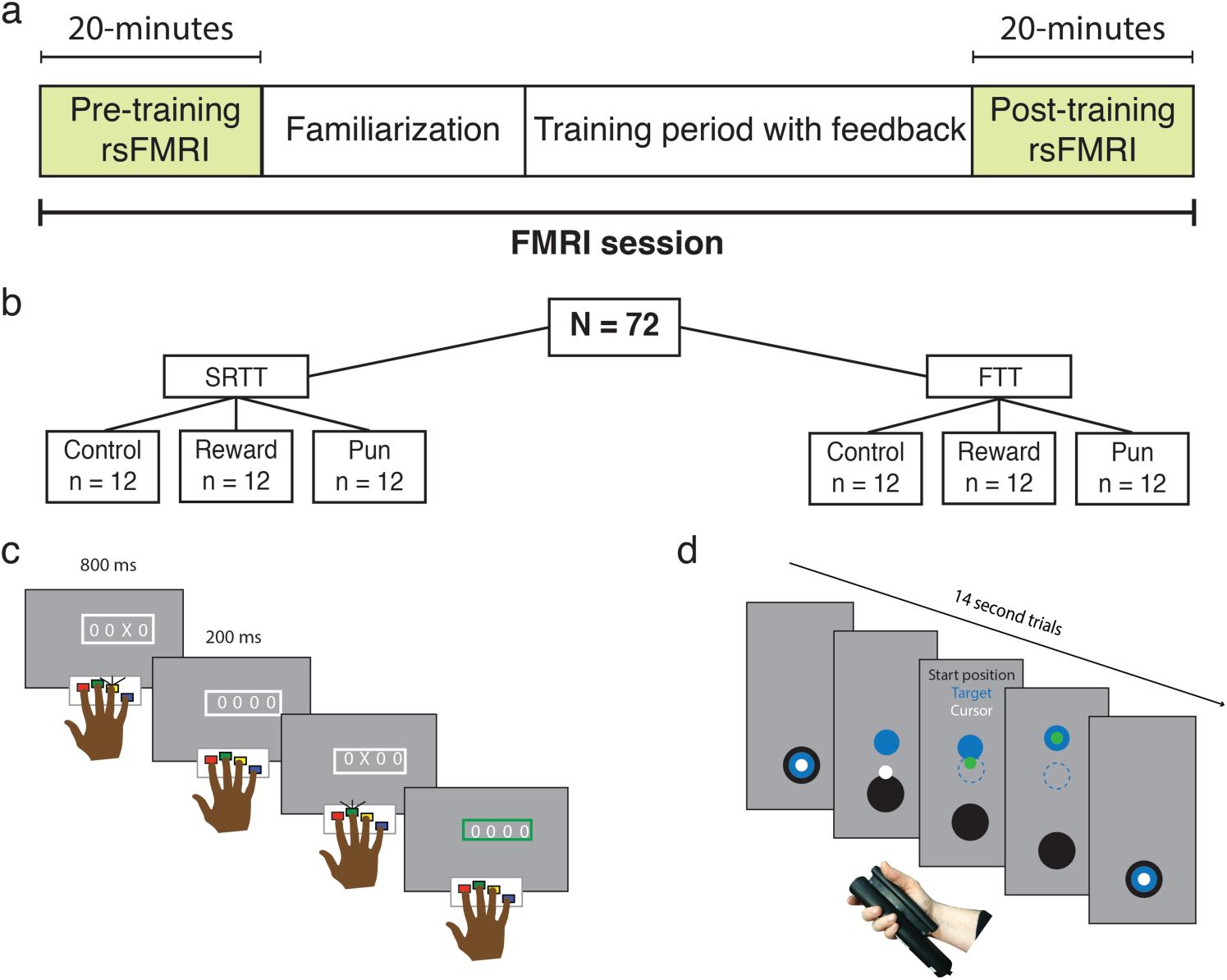
Experimental design and skill memory retention. (a,b) Participants underwent 20 minutes of resting state fMRI before and after training on either the serial reaction time task (SRTT) or the force tracking task (FTT) while receiving reward, punishment, or control feedback. In the SRTT (c), participants responded to a cue appearing in one of four locations on a screen. In the FTT (d), participants modulated their grip force to track a moving target. In both tasks, the stimulus could follow either a random- or fixed-sequence, and skill memory was assessed by comparing performance during random- and fixed-sequence trials.

We focused on premotor cortex (PMC) as the key region for evaluating post-encoding connectivity in both tasks, based on its well-documented critical role as a memory-encoding region for sequence and sensorimotor learning (Floyer-Lea and Matthews, 2004, 2005; Hardwick et al., 2013b; Kornysheva and Diedrichsen, 2014; Wiestler and Diedrichsen, 2013). In addition, PMC shows reward-related activity after movement (Ramkumar et al., 2016). Based on prior work (Murty et al., 2012a; Murty et al., 2016; Peterson and Seger, 2013a; Wachter et al., 2009), we hypothesized that connectivity between the PMC and the anterior insula, MTL, cerebellum, and caudate provide distinct contributions to skill learning.

## Materials and methods

### Overview

In two experiments, participants were trained on either the serial reaction time task (SRTT) or the force-tracking task (FTT) with reward, punishment, or uninformative feedback (Figure 1b-d). No participant was trained in both tasks. A detailed description of the tasks and training procedure can be found in (Steel et al., 2016). Before and after the training session, 20-minutes of resting-state fMRI was collected.

### Participants

78 participants (47 female, mean age = 25 years ± std. 4.25) were recruited and participated in the study. All participants were right-handed, free from neurological disorders, and had normal or corrected-to-normal vision. All participants gave informed consent and the study was performed with National Institutes of Health Institutional Review Board approval in accordance with the Declaration of Helsinki (93-M-0170, NCT00001360). Data from 6 individuals (2 female) were removed from the study due to inattention during training (defined as non-responsive or inaccurate performance on greater than 50% of trials; n=3) or inability to complete the imaging session due to discomfort or fatigue (n=3). This left 72 participants with complete data sets included in the analyses presented here.

### Training procedure

Both tasks followed the same behavioral training procedure. Trials were presented over 15 blocks with a 30-second break separating each. Unbeknownst to the participants, during some blocks (“fixed-sequence blocks”) the stimulus would appear according to a repeating pattern (described below for each task). During other blocks the appearance of the stimulus was randomly determined (“random-sequence blocks”).

To familiarize participants to the task, and establish their baseline level of performance, the task began with three random-sequence blocks without feedback (“familiarization blocks”). Participants were unaware of the forthcoming feedback manipulation during the familiarization blocks. Then the feedback period began, starting with a pre-training probe (three blocks, random – fixed – random), then the training blocks (six consecutive fixed-sequence blocks), and, finally, a post-training probe (three blocks, random – fixed – random). To test the impact of reward and punishment on skill learning, participants were randomized into one of three feedback groups: reward, punishment, or uninformative (control). The feedback paradigm for each task is outlined separately below.

Training was conducted inside the MRI scanner, and fMRI data were collected during the training period. These training period data are outside the scope of the present manuscript and are not presented here.

### Experiment 1: Serial reaction time task (SRTT)

The version of the SRTT used here adds feedback to the traditional implementation. At the beginning of each block participants were presented with four “O”s, arranged in a line at the center of the screen. These stimuli were presented in white on a grey background (Figure 1c). A trial began when one of the “O”s changed to an “X”. Participants were instructed to respond as quickly and accurately as possible, using the corresponding button, on a four-button response device held in their right hand. The “X” remained on screen for 800 ms regardless of whether the subject made a response, followed by a 200 ms fixed inter-trial interval, during which time the four “O”s were displayed. The trial timing used in this study may foster some degree of explicit awareness, and therefore this variation of the SRTT should not be considered a pure motor learning task. However, this timing was necessary to accommodate the constraints of fMRI scanning and does not impact the analysis of the data presented herein.

In the SRTT, each block consisted of 96 trials. During fixed-sequence blocks, the stimuli appeared according to one-of-four fixed 12-item sequences, which repeated 8 times (e.g. 3-4-1-2-3-1-4-3-2-4-2-1). For each participant, the same 12-item sequence was used for the duration of the experiment. Each fixed-sequence block began at a unique position within the sequence, to help prevent explicit knowledge of the sequence from developing (Schendan et al., 2003). In the random-sequence blocks, the stimuli appeared according to a randomly generated sequence, without repeats on back-to-back trials, so, for example, subjects would never see the triplet 1-1-2.

Breaks between blocks lasted 30-seconds. Initially, participants saw the phrase “Nice job, take a breather”. After five seconds, a black fixation-cross appeared on the screen. Five seconds before the next block began, the cross turned blue to alert the subjects that the next block was about to start.

During the post-training retention probes, participants performed three blocks (random – fixed – random), outside the scanner on a 15-inch Macbook Pro using a button box identical to the one used during training. During these retention probes, the next trial began 200 ms after the participant initiated their response rather than after a fixed 800 ms as during training. No feedback was given during the retention blocks.

### Experiment 2: Force-tracking task

In the force-tracking task (FTT), participants continuously modulated their grip force to match a target force output (Floyer-Lea and Matthews, 2005; Floyer-Lea et al., 2006). In the traditional implementation, participants are exposed to a single pattern of force modulation that is repeated on every trial. This design does not allow discrimination between general improvement (i.e. familiarization with the task and/or the force transducer) and improvement specific to the trained sequence of force modulation. Therefore, we adapted the traditional FTT method to align it with the experimental design that is traditional for the SRTT, i.e. by including random-sequence blocks.

A given trial consisted of a 14 s continuous pattern of grip modulation. At the beginning of a trial, participants were presented with three circles on a grey background: a white circle (Cursor, 0.5 cm diameter), a blue circle (Target, 1.5 cm diameter), and a black circle (Bottom of the screen, 2 cm diameter; Figure 1d). Participants held the force transducer (Current Designs, Inc., Philadelphia, PA) in the right hand between the four fingers and palm. Application of force to the transducer caused the cursor to move vertically on the screen. Participants were instructed to keep the cursor as close to the center of the target as possible as the target moved.

For fixed-sequence blocks, participants were assigned to one of six possible 14 s patterns of grip modulation. This pattern was repeated on every sequence trial. During random-sequence blocks, the target followed a trajectory generated by the linear combination of four waveforms, with periods between 0.01 and 3 Hz. The combinations of waveforms were constrained to have identical average amplitude (target height), and the number and value of local maxima and minima were constant across the random blocks.

For data analysis, the squared distance from the cursor to the target was calculated at each frame refresh (60 Hz). The first 10 frames were removed from each trial. The mean of the remaining time points was calculated to determine performance, and trials were averaged across blocks.

### Feedback

All participants were paid a base remuneration of $80 for participating in the study. At the start of the feedback period, participants were informed they could earn additional money based on their performance.

In the SRTT, performance was defined as the accuracy (correct or incorrect) and reaction time (RT) of a given trial. Feedback was given on a trial-by-trial basis (Figure 1c,d). This was indicated to the participant when the white frame around the stimulus changed to green (reward) or red (punishment). Participants in the reward group were given feedback if their response was accurate and their RT was faster than their criterion RT, which indicated that they earned money ($0.05 from a starting point of $0) on that trial. Participants in the punishment group were given feedback if they were incorrect, or their RT was slower than their criterion, which indicated that they lost money ($0.05 deducted from a starting point of $55) on that trial. Participants in the control-reward and control-punishment groups saw red or green color changes, at a frequency matched to punishment and reward; the color changes were not related to their performance. Control participants were told that they would be paid based on their speed and accuracy.

Feedback in the FTT was based on the distance of the cursor from the target. For the reward group, participants began with $0. As participants performed the task, their cursor turned from white to green whenever the distance from the target was less than their criterion, indicating that they were gaining money at that time. Participants in the punishment group began with $45, and, as they performed, the cursor turned red if it was outside their criterion distance. This indicated that they were losing money. For reward-control and punishment-control groups, the cursor changed to green or red. As in the SRTT, the color changes for the control group were not related to the participant’s performance. For control, the duration of each feedback instance, as well as cumulative feedback given on each trial, was matched to the appropriate group.

In both tasks, for the reward and punishment groups, between blocks, the current earning total was displayed (e.g. “You have earned $5.00”). Control participants saw the phrase, “You have earned money.” The criterion RT was calculated as median performance in the first familiarization block. After each block, the median + standard deviation of performance was calculated, and compared with the criterion. If this test criterion was faster (SRTT) or more accurate (FTT) than the previous criterion, the criterion was updated. During the SRTT, only the correct responses were considered when establishing the criterion reaction time.

Importantly, to control for the motivational differences between gain and loss, participants were not told the precise value of a given trial. This allowed us to assess the hedonic value of the feedback, rather than the level on a perceived-value function.

### MRI acquisition

This experiment was performed on a 3.0T GE 750 MRI scanner using a 32-channel head coil (GE Medical Systems, Milwaukee, WI).

### Structural scan

For registration purposes, a T1-weighted anatomical image was acquired (magnetization-prepared rapid gradient echo (MPRAGE), TR = 7 ms, TE = 3.4 ms, flip-angle = 7 degrees, bandwidth = 25.000 kHz, FOV = 24×24 cm^2^, acquisition matrix = 256 × 256, resolution = 1 × 1 × 1 mm, 198 slices per volume). Grey matter, white matter, and CSF maps for each participant were generated using Freesurfer (Fischl et al., 2002).

### EPI scans

Multi-echo EPI scans were collected with the following parameters: TE = 14.9, 28.4, 41.9 ms, TR = 2, ASSET acceleration factor = 2, flip-angle = 65 degrees, bandwidth = 250.000 kHz, FOV = 24 × 24 cm, acquisition matrix = 64 × 64, resolution = 3.4 × 3.4 × 3.4 mm, slice gap = 0.3 mm, 34 slices per volume covering the whole brain. Respiratory and cardiac traces were recorded. Each resting state scan lasted 21-minutes. The first 30 volumes of each resting-state scan were discarded to control for the difference in arousal that occurs at the beginning of resting state scans. This left the final 20-minutes of rest in each scan for our analysis. This procedure has been used in other studies where long-duration resting state runs were collected (Gonzalez-Castillo et al., 2014).

### Resting state fMRI preprocessing

Data were preprocessed using AFNI (Cox, 1996). The time series for each TE was processed independently prior to optimal combination (see below). Slice-time correction was applied (3dTShift) and signal outliers were attenuated [3dDespike (Jo et al., 2013)]. Motion correction parameters were estimated relative to the first volume of the middle TE (28.4 msec), and registered to the structural scan (3dSkullStrip, 3dAllineate). These registration parameters were then applied in one step (3dAllineate) and the data were resampled to 3 mm isotropic resolution.

The optimal echo time for imaging the BOLD effect is where the TE is equal to T2*. Because T2* varies across the brain, single echo images are not optimal to see this variation. By acquiring multiple echoes, we could calculate the “optimal” T2* value for each voxel through the weighted average of the echoes at each time point, which allowed us to recover signals in dropout areas and improves contrast-to-noise ratio (Evans et al., 2015; Kundu et al., 2014; Poser et al., 2006; Posse et al., 1999). The following is a summary of methods implemented for optimal combination implemented in meica.py [(Kundu et al., 2012)].

The signal at an echo, *n* varies as a function of the initial signal intensity *S_0_* and the transverse susceptibility T2* = 1/R2* and is given by the mono-tonic exponential decay:

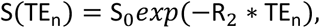
where R2* is the inverse of relaxation time or 1/T2*. This equation can be linearized to simplify estimation of T2* and S_0_ as the slope using log-linear transformation. The time courses can be optimally combined by weighted summation by a factor, w, described by the following equation:

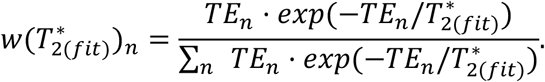

Where T_2_(fit) is the transverse relaxation time estimated for each voxel using the equation above. The optimally combined time series can then be treated as a single echo.

After optimal combination, we applied the basic ANATICOR (Jo et al., 2010) procedure to yield nuisance time series for the ventricles and local estimates noise from the white matter. All nuisance time-series (six parameters of motion, local white matter, ventricle signal, and 6 physiological noise regressors (RetroTS)) were detrended with fourth order polynomials. These time series, along with a series of sine and cosine functions to remove all frequencies outside the range (0.01–0.25 Hz) were projected out of the data in a single regression step (3dTproject). Time points with motion greater than 0.3 mm were removed from the data [scrubbing, see Power et al. (2012)] and replaced with values obtained via linear interpolation in time. Data were aligned to the N27 atlas and transformed into Talairach space (@auto_tlrc) and smoothed with a 6mm FWHM Gaussian kernel. For group data analysis, a group-level grey matter by mask was created by calculating voxels determined to be grey matter in 80% of participants (Gotts et al., 2012). Global signal regression was not performed on these data (Saad et al., 2012).

### Left premotor cortex functional connectivity

We focused our analysis on the PMC based on this region’s well-described central role in sequence production in response to visual cues (Mushiake et al., 1991), sensorimotor learning, and sequence learning (Hardwick et al., 2015; Hardwick et al., 2013b). Given that the participants were performing the task with their right hand, we further focused on the left PMC. Left dorsal and ventral PMC (PMd/PMv) were defined based on a publicly available diffusion-MRI-based parcellation of premotor cortex (Tomassini et al., 2007). The PMd and PMv regions of this atlas are matched for size.

The mean time series from both dorsal and ventral premotor cortices were extracted separately from each participant and each rest period. The whole brain correlation maps (Pearson’s r) for both PMd and PMv during the pre- and post-training resting state MRI scans were then calculated based on these time series. Prior to running group statistics, these maps were Z-scored using a Fisher’s transform. For each resting-state scan, global correlation [GCOR; @compute_gcor: Saad et al. (2013)] and magnitude of motion across runs (@1dDiffMag) were calculated and included as nuisance covariates for group analysis.

The resulting maps were then submitted to a linear mixed effects model (3dLME) with ROI (PMd/PMv), Rest (pre-/post-), Feedback valence (Control/Reward/Punishment) as factors. The precise model fit at the group level in R-syntax (nlme) was ‘Feedback valence*Rest*ROI+motion+gcor’. A random effect (subject) was included in the model. Group analysis maps were cluster-corrected for multiple comparisons to achieve a α = 0.05 using the ACF model in 3dClustSim (p < 0.005, k = 54; AFNI compile date July 9, 2016).

To determine the consistency of the impact of feedback across the two tasks, the overlap of the significant clusters from the Rest x Feedback valence interaction for the SRTT and FTT was calculated at both a conservative (p < 0.005, k = 54) and liberal (p < 0.01, k = 54) threshold. In addition, to formally test for a difference between the feedback valence conditions across tasks, we considered both tasks together in a single model that included Task (SRTT/FTT) as a factor. The precise model fit for this analysis was ‘Task* Feedback valence*Rest*ROI+motion+gcor’.

## Results

The behavioral data have previously been reported [Steel et al. (2016)]. Briefly, during performance of the SRTT, we found that, compared to participants training with reward, those training with punishment showed reduced reaction time without detriment to accuracy; in contrast, on the FTT, compared to participants training with reward, those training with punishment exhibited greater tracking error during training (Supplemental Figure 1). During retention testing, in both tasks all feedback groups showed evidence of sequence knowledge at 1hr, 24–48hrs and 30-d after training. To identify brain regions impacted by feedback given during training, we used a seed-based analysis focused on the left PMC. For each experiment, we implemented a voxel-wise linear mixed effects (LME) model (Chen et al., 2013) with Rest (Pre-/Post-training), Feedback valence (Reward /Punishment /Control), and ROI (ventral-/dorsal-premotor cortex) as factors and looked for regions showing a significant interaction between Rest x Feedback valence. No regions showed a Rest x Feedback valence x ROI interaction in either task. Full results from this model for both tasks can be found in Table 1.

**Table 1.** Significant clusters for SRTT and FTT voxel-wise linear mixed effects model analyses. Group analysis maps were cluster-corrected for multiple comparisons to achieve a α = 0.05 using the ACF model in 3dClustSim (p < 0.005, k = 54; AFNI compile date July 9, 2016).

| **Number of voxels** | **Volume** | **x** | **y** | **z** | **Region** |
| --- | --- | --- | --- | --- | --- |
| **SRTT** |  |  |  |  |  |
| *Main effect Rest* |  |  |  |  |  |
| 1909 | 51543 | -4.5 | -22.5 | 11.5 | Left Thalamus |
| 949 | 25623 | 52.5 | 1.5 | -6.5 | Right superior temporal gyrus |
| 477 | 12879 | 1.5 | -73.5 | -24.5 | Right cerebellar vermis |
| 299 | 8073 | -55.5 | -1.5 | -3.5 | Left superior temporal gyrus |
| 68 | 1836 | 25.5 | -13.5 | -12.5 | Right parahippocampal cortex |
| *Rest x Condition* |  |  |  |  |  |
| 828 | 22356 | -16.5 | 1.5 | -0.5 | Left pallidum |
| 215 | 5805 | 28.5 | 1.5 | -18.5 | Right parahippocampal cortex |
| 195 | 5265 | -10.5 | -34.5 | 59.5 | Left supplementary motor area |
| 154 | 4158 | -4.5 | -37.5 | -24.5 | Left cerebellum |
| 77 | 2079 | -16.5 | -13.5 | -18.5 | Left parahippocampal cortex |
| 59 | 1593 | -31.5 | 19.5 | 15.5 | Left inferior frontal gyrus |
| **FTT** |  |  |  |  |  |
| *Main effect Rest* |  |  |  |  |  |
| 3545 | 95715 | -4.5 | -22.5 | 11.5 | Left thalamus |
| 148 | 3996 | 7.5 | -31.5 | 59.5 | Right supplementary motor area |
| 118 | 3186 | -1.5 | 55.5 | -6.5 | Left medial frontal gyrus |
| 112 | 3024 | 43.5 | -61.5 | -3.5 | Left middle occipital gyrus |
| 112 | 3024 | 1.5 | -61.5 | 17.5 | Right posterior cingulate cortex |
| 60 | 1620 | 16.5 | -73.5 | 29.5 | Right precuneus |
| *Rest x Condition* |  |  |  |  |  |
| 639 | 17253 | 46.5 | -49.5 | -15.5 | Right fusiform gyrus |
| 395 | 10665 | -49.5 | -64.5 | 8.5 | Left middle temporal gyrus |
| 147 | 3969 | -10.5 | -91.5 | 5.5 | Left cuneus |
| 124 | 3348 | 25.5 | -73.5 | 38.5 | Right precuneus |
| 74 | 1998 | 13.5 | 1.5 | 5.5 | Right pallidum |
| 63 | 1701 | 19.5 | -61.5 | -21.5 | Right cerebellum lobule VI |
| 60 | 1620 | -7.5 | -64.5 | 2.5 | Left lingual gyrus |
| 55 | 1485 | 7.5 | 61.5 | 2.5 | Right medial frontal gyrus |

### Reward and punishment evoke dissociable changes in PMC connectivity patterns after SRTT training

In Experiment 1, following training on the SRTT task, PMC functional connectivity change due to training was modulated by feedback valence in several regions (Rest x Feedback valence interaction; Figure 2). These regions included bilateral thalamus and striatum, right cerebellar vermis, supplementary motor area (SMA), bilateral medial temporal lobe, and left inferior frontal gyrus. In order to examine the nature of the interaction, we extracted the estimated mean PMC functional connectivity change due to training from each cluster (Figure 2, inset). This revealed a pattern that was distinct across the feedback groups. After training with reward, functional connectivity increased between PMC and thalamus and striatum, cerebellar vermis, and SMA, but PMC functional connectivity decreased with MTL and inferior frontal gyrus. The punishment group showed the opposite pattern; PMC connectivity with medial temporal lobe and inferior frontal gyrus increased after training with punishment. The control group showed an intermediate pattern between the reward and punishment groups. In the control group, PMC connectivity increased with the thalamus, striatum, and cerebellar vermis after training, but decreased with SMA, medial temporal lobe, and inferior frontal gyrus. Thus, in the context of the SRTT, reward and punishment have clearly differentiable impacts on functional connectivity change due to training in regions associated with learning on this task.

**Figure 2.**
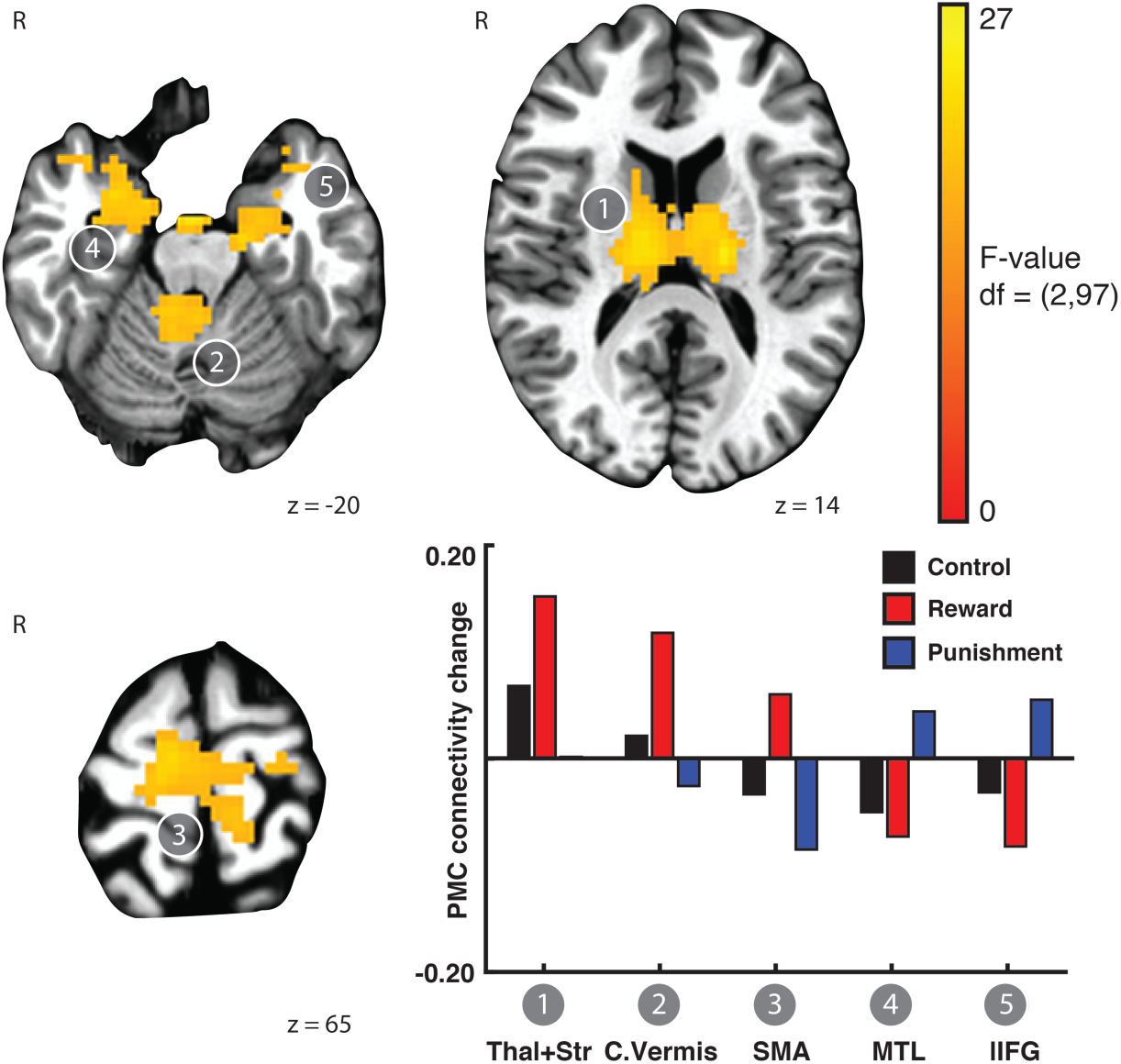
Reward and punishment differentially affect PMC functional connectivity change after training on the SRTT. Linear mixed effects modelling revealed brain regions that exhibited a Rest x Feedback valence interaction in the functional connectivity of left PMC. Functional connectivity between PMC and the thalamus and striatum, cerebellum, and SMA increased after training with reward and control feedback but decreased after training with punishment. In contrast, functional connectivity between medial temporal lobe and left inferior frontal gyrus increased after training with punishment but decreased after training with reward and control feedback. Bars showing mean connectivity change across voxels in the identified cluster after training estimated by the linear mixed effects model are included to enable qualitative comparison and reveal the nature of the interaction effect.

### Punishment promotes PMC-striatal connectivity after FTT training

In Experiment 2, after training on the FTT, PMC connectivity change was also affected by feedback valence (LME: Rest x Feedback valence interaction). However, the regions exhibiting the Rest x Feedback valence interaction after training on the FTT differed from those found in after training on the SRTT (Experiment 1). After training on the FTT, we found that PMC connectivity with bilateral lateral occipital temporal cortex, precuneus, left culmen, cerebellum, right striatum, and right medial frontal gyrus differed across the Feedback valence conditions (Figure 3). Extracting the estimated mean PMC functional connectivity change of these regions revealed that two primary modes of change (Figure 3, inset). PMC connectivity with the lateral occipital cortex, precuneus, culmen, and cerebellar lobule VI, decreased after training with punishment, but increased after training with control feedback. In contrast, connectivity of the PMC with right striatum and right medial frontal gyrus increased after training with punishment. PMC connectivity to right medial frontal gyrus decreased after training with control. Overall, in the regions showing the interaction between functional connectivity change after training and feedback valence, reward had a smaller influence compared to control or punishment. This suggests that punishment induces marked changes in functional connectivity change after FTT training, which was consistent with the behavioral effects observed at the end of training (Steel et al., 2016).

**Figure 3.**
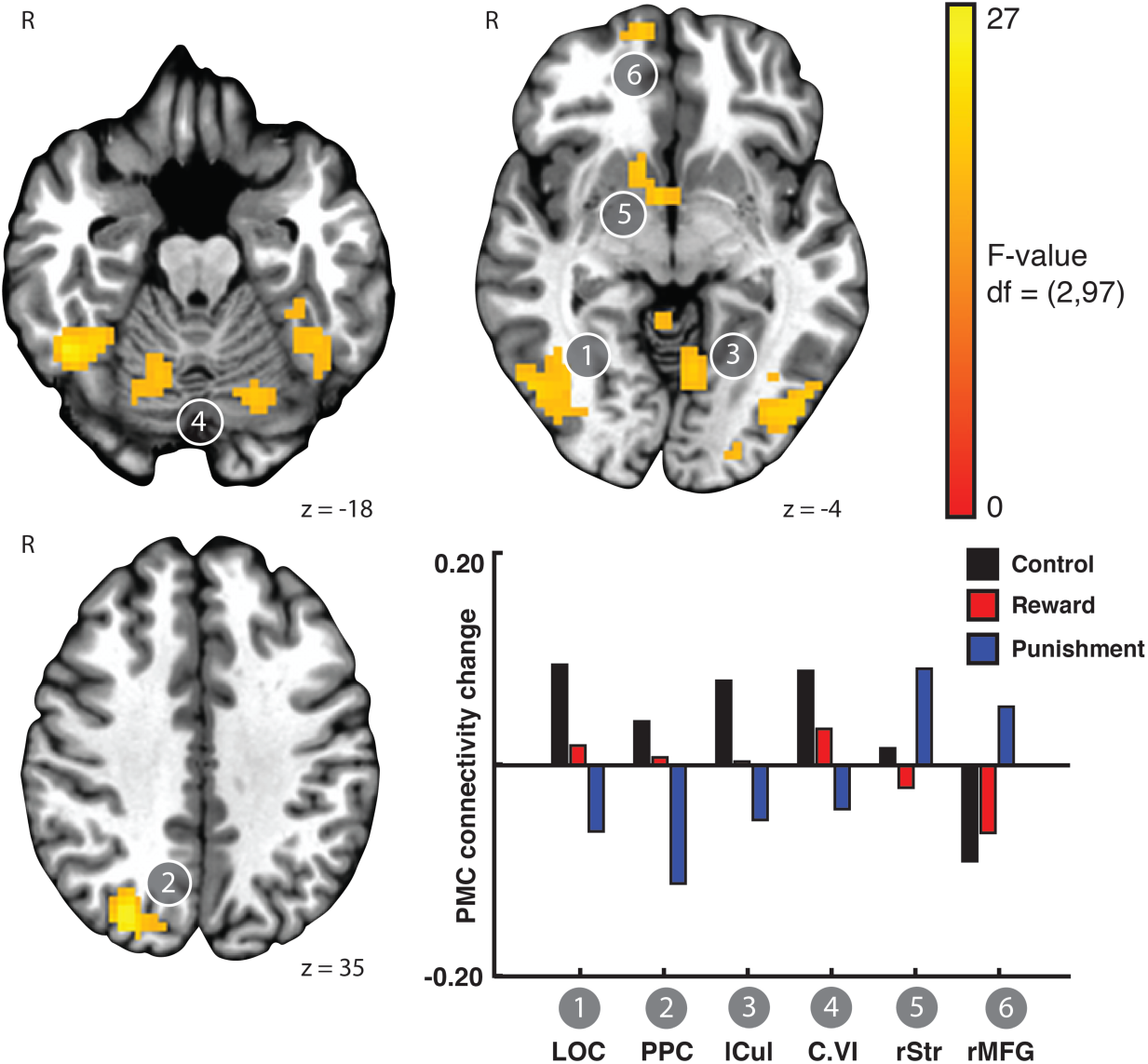
Punishment promotes PMC-striatal connectivity after FTT training. A linear mixed effects model revealed brain regions exhibiting a Rest x Feedback valence interaction in the functional connectivity of left PMC. Functional connectivity between the PMC and lateral occipital cortex, posterior parietal cortex, culmen, and cerebellum increased after training with control feedback, but decreased after training with punishment. In contrast, PMC connectivity with right ventral striatum and right medial frontal gyrus increased after training with punishment but decreased after training with reward and control feedback. Bars showing mean connectivity change across voxels in the identified cluster after training estimated by the linear mixed effects model are included to enable qualitative comparison.

### Influence of reward and punishment on PMC functional connectivity depends on task

In order to better understand the correspondence between feedback valence and the change in PMC functional connectivity after training between the two tasks, we calculated the overlap between the sets of regions that showed the Rest x Feedback valence interaction for each task separately. Given the reduced power to detect general effects that might be weak but consistent across tasks, for this qualitative assessment we considered the conservative threshold reported above (p < 0.005, k = 54), as well as a liberal threshold (p < 0.01, k = 54; Figure 4a). The mean connectivity change with PMC for clusters greater than 20 voxels showing an overlap between the two tasks is shown in Figure 4b.

**Figure 4.**
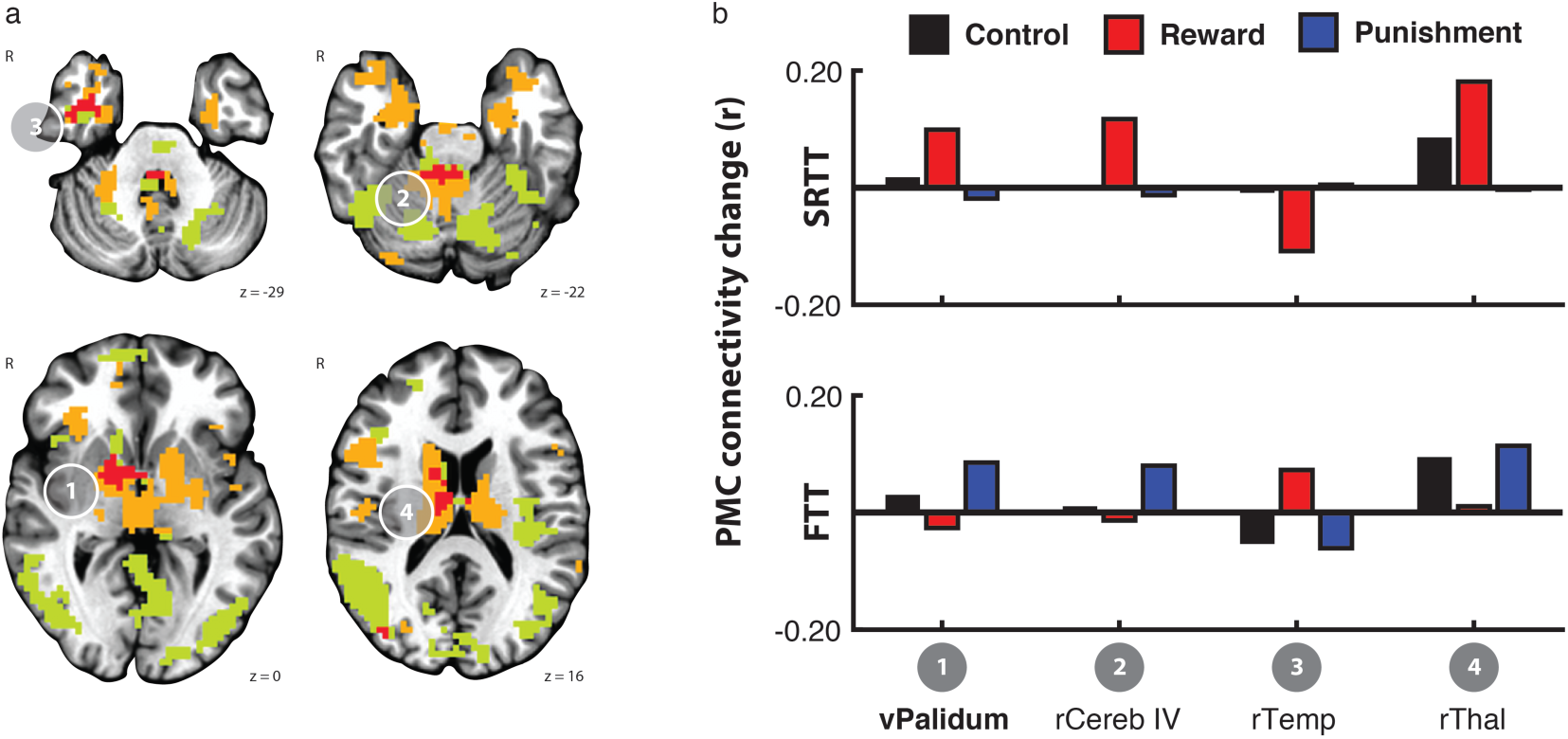
Regions showing PMC functional connectivity change after training had limited overlap across the two tasks. Clusters that showed a PMC connectivity change after training that differed by the feedback group in the SRTT (orange) and the FTT (green) showed limited overlap, even at a liberal cluster threshold (p < 0.01, k = 54). Only the ventral pallidum (cluster 1, bold) showed an overlap at the threshold reported above (p < 0.005, k = 54). In all clusters, the reward and punishment groups exhibited opposite patterns of connectivity change between PMC and the overlapping regions due to training on the SRTT (upper) and FTT (lower). Bars showing mean connectivity change across voxels in the identified cluster after training estimated by the linear mixed effects model are included to enable qualitative comparison.

At the conservative threshold, one region in the striatum showed a Rest x Feedback valence interaction in both tasks (Cluster 1). When we extracted the estimated mean connectivity change after training for each task, however, we found that the pattern of connectivity change across the feedback valence groups was diametrically opposed. In the SRTT, connectivity between the PMC and the overlapping region increased after training with reward but decreased after training with punishment. In contrast, after training on the FTT, connectivity between PMC and the overlapping region increased after training with punishment but increased after training with reward. In both tasks, connectivity between PMC and the overlapping region increased after training with control feedback.

At the liberal threshold, we detected three other clusters in the right anterior temporal lobe, the right cerebellum, and right dorsal thalamus. In each of these clusters, the pattern of connectivity change with PMC exhibited by the reward and punishment groups was also not consistent across tasks.

To directly compare the effect of task on the functional connectivity change after training with feedback, we performed a whole-brain analysis (cluster corrected; p < 0.005, k = 54) using a linear mixed effects model that included Task (SRTT/FTT), Rest (Pre-/Post-training), and Feedback valence (Control/Reward/Punishment) as factors. For this analysis, we specifically focused on regions that showed a Task x Feedback valence x Rest interaction. All regions that appeared in the overlap analysis reported above evidenced a Task x Feedback valence x Rest interaction (bilateral basal ganglia including caudate, putamen, and pallidum [peak x=-10, y=16, z=17], right cerebellum [1.5, 24.5, -24.5], right medial temporal lobe [-40, 4.5, -33.5]). In addition, the right lateral temporal lobe [-43, 49, -15], right parietal cortex [-43, 55, 17.5], right medial frontal gyrus [-13.5, 52.5, -6.5], and right lateral prefrontal cortex [-46.5 -16.5 17.5] also showed this interaction (Figure 5).

**Figure 5.**
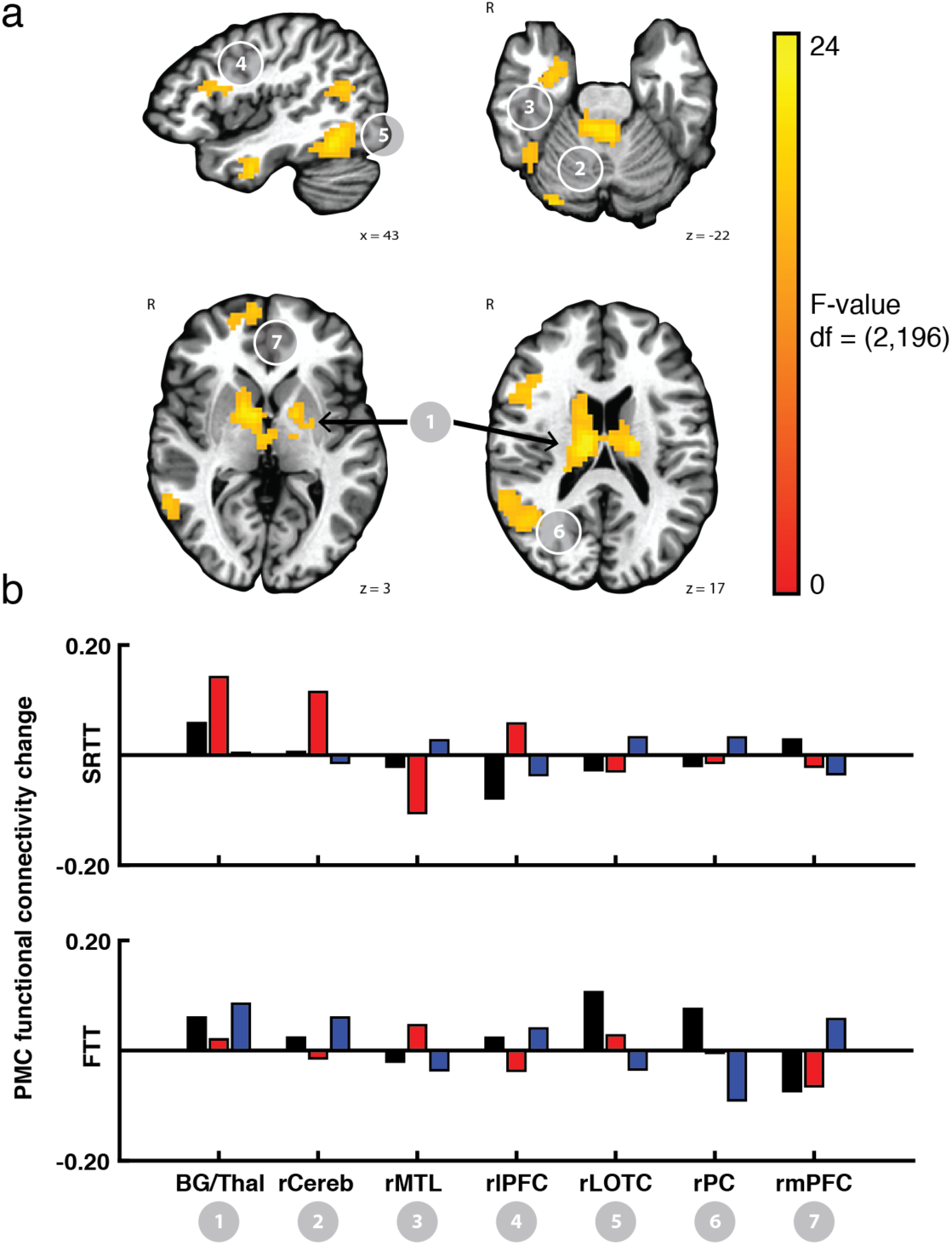
Difference in PMC functional connectivity due to training across the feedback valence groups depends on the task being performed. Upper. Data from both tasks were submitted to a linear mixed effects model. This analysis revealed the resulting F-statistic map for the interaction term ‘Task*Feedback valence*Rest’ indicating regions within the basal ganglia, cerebellum, MTL, temporal cortex, parietal cortex, and prefrontal cortex showed an effect of feedback on functional connectivity change after training that varied by task. Mean PMC functional connectivity change after training from these ROIs is shown in the lower panel. Importantly, all ROIs that showed an overlap in Figure 4 also showed the interaction in the LME analysis. Bars showing mean connectivity change across voxels in the identified cluster after training estimated by the linear mixed effects model are included to enable qualitative comparison.

In summary, it is clear that feedback valence profoundly impacts resting state functional connectivity changes after training, but that the regions of the brain impacted by feedback valence depend on the task being performed.

## Discussion

In this experiment, we sought to investigate the effect of feedback valence on brain activity during the period immediately SRTT and FTT training. We found that reward and punishment differentially impacted change in PMC functional connectivity, and this impact was influenced by task. After training on the SRTT, functional connectivity between PMC with striatum, cerebe0llum, and SMA increased after training with reward, while functional connectivity between PMC with medial temporal lobe and inferior frontal gyrus increased after training with punishment. Training on the FTT induced a different pattern of connectivity change across the feedback valence groups. After FTT training, functional connectivity between PMC and parietal cortices increased after training with control feedback. In contrast, functional connectivity between PMC and parietal cortices decreased after training with punishment, while functional connectivity between PMC with the striatum and medial frontal gyrus increased after training with punishment. Finally, we looked for regions that showed an overlap in the Rest x Feedback valence interaction across tasks. At our conservative threshold, the clusters of significant voxels for the two tasks overlapped only in the ventral pallidum. In the overlapping voxels, the two tasks showed opposite patterns of results: after training on the SRTT, connectivity between the PMC and the pallidum increased after training with reward but decreased after training with punishment; the opposite pattern was true for the FTT, after which connectivity between the PMC and the pallidum increased after training with punishment but decreased after training with reward. Taken together, these results suggest that training with valenced feedback has differential effects on functional connectivity after training, and that these effects primarily manifest in the regions involved in task performance.

Reward and punishment differentially impacted PMC functional connectivity with the striatum and medial temporal lobe after training on the SRTT. SRTT learning engages the motor system, including motor cortex and premotor cortex (Hardwick et al., 2013b; Keele et al., 2003; King et al., 2017; Kornysheva and Diedrichsen, 2014; Schubotz and von Cramon, 2003; Wiestler and Diedrichsen, 2013), parietal cortex (Breton and Robertson, 2017; Keele et al., 2003), basal ganglia (Albouy et al., 2013a; Carbon et al., 2004; Debas et al., 2014; Seger, 2006), and medial temporal lobe (Albouy et al., 2013b; Schendan et al., 2003). These structures are often classified as fast (frontoparietal and hippocampal) and slow (motor and basal ganglia) systems, based on the onset of evoked activity of these regions during the learning process (Doyon et al., 2018). One theory to explain the impact of reward and punishment on memory formation is the engagement of the slow and fast systems, respectively. With respect to the SRTT, we found evidence for this differential recruitment of the fast and slow learning systems after training with punishment and reward, respectively. PMC connectivity with motor and basal ganglia structures increased after training with reward and PMC connectivity with anterior insula and hippocampus after increased training with punishment.

In addition to reward and punishment, other factors could interact with brain areas recruited during SRTT learning. For example, the contribution of the motor, parietal, and subcortical systems may be based on the goal of the learner, or based on the statistical complexity of the sequence being learned (Robertson, 2007). However, it is clear that many brain regions interact during the learning process. For example, it is possible to bias neural activity towards one set of brain areas [e.g. (Keele et al., 2003)] or to induce competitive interactions among the memory systems by constraining task conditions (Brown and Robertson, 2007; Tunovic et al., 2014). There has been evidence to suggest that valenced feedback could also dissociate the brain areas activated during skill-learning (Wachter et al., 2009), and the present data extend this knowledge by showing that reward and punishment can bias recruitment of neural areas after learning.

Reward and punishment also had differential effects on functional connectivity after FTT training, but in different regions from the SRTT. After training on the FTT, we observed differences amongst the feedback valence groups in functional connectivity between PMC and posterior parietal, lateral occipital, and prefrontal cortices, as well as the ventral striatum and the cerebellum. The control and reward groups showed increased connectivity between PMC and the parietal, occipital, and cerebellar regions, but decreased connectivity between PMC and ventral striatum and medial frontal gyrus. The punishment group showed the opposite pattern: functional connectivity between PMC and parietal, occipital, and cerebellum decreased after training and PMC connectivity with the striatum and medial frontal gyrus increased after training.

FTT learning is generally associated with decreased BOLD activity in the cortical motor network, prefrontal cortex, and caudate nucleus, but increased BOLD activity in the cerebellum and putamen (Floyer-Lea and Matthews, 2004, 2005). While prior studies of FTT training did not find BOLD-signal activity changes in lateral occipital cortex, this region is strongly associated with action representation (Kable and Chatterjee, 2006; Kable et al., 2005; Lingnau and Downing, 2015) and may be consistently engaged while learning the novel visuomotor association for learning on the FTT. Thus, the effects of feedback valence are largely restricted to areas involved in performance of the task.

Overall with respect to the influence of feedback valence on functional connectivity change due to training, we observed very little overlap between the FTT and SRTT. At the conservative threshold, the only region where feedback modulated PMC functional connectivity change due to training on both tasks was the ventral pallidum. The center of mass of the cluster overlap was centered in the medial pallidum, and the cluster extent encompassed both the lateral and medial pallidum. Across the two tasks, reward and punishment had opposing effects on PMC connectivity with this area. After training on the SRTT, functional connectivity between the PMC and ventral pallidum increased after training with reward and decreased after training with punishment. The opposite was true after training on the FTT, wherein we observed increased functional connectivity between PMC and the ventral pallidum after training with punishment, but a decrease after training with reward. The ventral pallidum contains cell populations that respond to both appetitive and aversive stimuli (Saga et al., 2017). Behavioral response to appetitive and aversive stimuli are mediated by glutamatergic and GABAergic neurons in the ventral pallidum, and project to the ventral tegmental area and lateral habenula, respectively (Faget et al., 2018). Therefore, given that cells in this area respond to positive and negative stimuli, it may not surprising to find this region affected by both reward and punishment following training in our study.

Across both tasks, the dissociation between the feedback valence groups with respect to PMC-pallidum functional connectivity change after training matched the dissociation in behavioral performance: the group that showed PMC-pallium functional connectivity increased after training performed worse on the task, while the group that showed diminished PMC-pallidum functional connectivity performed better at the end of training [see Supplemental Figure 1 for a summary of the relevant behavioral data and (Steel et al., 2016) for behavioral results of the participants in this study]. Specifically, on the SRTT, the punishment group performed better during training (i.e. had a faster overall reaction time) compared to the reward and control groups, and, on this task we observed a decrease in functional connectivity between ventral pallidum and PMC after training with punishment. On the other hand, in the FTT, the punishment group performed worse (i.e. made more tracking error) than the reward and control groups behaviorally, and the connectivity between PMC and ventral pallidum increased after training on the FTT with punishment. Thus, with respect to the pallidum, it may be that the connectivity of this region to PMC depends on the experience of the learner (either absence of reward or presence of punishment) in the environment, and this activity may persist after training.

A meta-analysis of decision-making studies using valenced feedback found surprise, valence, and signed prediction error information converged specifically in the ventral pallidum (Fouragnan et al., 2018). It is notable that the pallidum itself is a heterogenous structure embedded within the corticobasal-ganglia thalamo-cortical loops (Redgrave et al., 2010). Even though this structure may play a common role integrating the error-related information in both tasks, this error-related information may be originating from distinct cortical networks projecting to separate aspects of the pallidum [e.g., associative versus motor (Redgrave et al., 2010)]. The scanning resolution used here only allows coarse localization of the effects within the pallidum, but this could be addressed using higher-resolution imaging modalities.

Even at a liberal threshold, the regions that showed overlap across the two tasks showed opposite patterns of connectivity change based on the type of feedback given. Similar to the pallidum, the differential effect of reward and punishment on functional connectivity between PMC and these regions may be due to the qualitative differences in performance. More generally, the minimal overlap and opposing patterns suggest that reward and punishment have strong impacts on post-training brain activity and that this effect is strongly influenced by the task that was performed.

Valenced feedback processing is often characterized as being mediated by a set of regions that would be common across tasks; for example, in the context of statistical learning and decision making, distinct networks for valence, surprise, and signed-prediction error have been suggested (Fouragnan et al., 2018). Regions activated in response to surprise and valence information include the striatum and insular cortex, which differentially responded to positive and negative surprises, respectively (Fouragnan et al., 2018). The regions showing changes in functional connectivity after training on the SRTT task match these predictions, and others have suggested that the SRTT is a statistical learning task (Robertson, 2007). After training on the FTT, the patterns of connectivity change across the feedback valence groups was not consistent with what would be expected after statistical learning. After training on FTT, we observed that task-relevant regions, including the lateral occipital, parietal, and cerebellar activity (Grafton et al., 2008; Hardwick et al., 2013b; Imamizu et al., 2000; Krakauer et al., 2004) showed feedback related changes in functional connectivity. There is some evidence that the cerebellum also plays a role in motivational aspects of movement correction during motor learning (Bostan and Strick, 2018; Turner and Desmurget, 2010; Turner et al., 2003), and it is likely that the engagement of cerebellum depends on task-demands, as well. In summary, our data provide further evidence that, rather than be restricted to feedback-processing networks that are segregated on the basis of valence, rewards and punishments may be processed in a distributed fashion among brain regions engaged by the task.

### Limitations

Several aspects of the present study are worth noting. First, we did not distinguish between dorsal and ventral premotor cortex in our analyses because no regions showed a Rest x Group x ROI interaction in either task. Dorsal and ventral premotor cortex are highly interconnected and although feedback valence may differentially impact dorsal and ventral premotor cortex, we may have been underpowered to detect any effect. Second, it should be noted that our implementation of the SRTT included a fixed trial length rather than a self-paced trial duration, which might foster explicit knowledge, and it has been shown that explicit learning recruits different neural networks after learning (Sami et al., 2014). No participants included in the study spontaneously reported sequence knowledge. When tested at 3+ weeks, participants showed no evidence of explicit awareness (for further discussion, see Steel et al., 2016a). In addition, in order to match the SRTT parameters, our force tracking task implementation featured continuous feedback of cursor position, rather than feedback at given at the end of the trial, and future work may consider the difference between continuous versus end-of-trial feedback. Third, it is widely known that sleep interacts with memory formation and offline memory processes (Albouy et al., 2013a; Robertson et al., 2004a; Robertson et al., 2004b). In our study, we did not measure sleep quality. Whether sleep interacts with the effect of reward and punishment on retention may be an interesting avenue for future research. Fourth, our analysis focused specifically on the impact of reward and punishment on PMC because of this region’s known importance to motor control and motor learning (Hardwick et al., 2013a). It is likely that reward and punishment impacts other regions not specifically located within the motor system. Future work might consider the impact of rewards and punishments on other areas of the brain, as well as during other types of tasks. Finally, we note that although we observed statistically significant changes in functional connectivity across the feedback valence groups, we cannot conclude that these changes solely reflect memory-formation related activity. In addition to memory-formation related activity, other factors such as task performance (rather than memory formation) and rumination also influence functional connectivity after training. Based on our experimental design and sample size, we cannot disambiguate these effects in our data. Follow-up studies may consider causal manipulations to assess the importance of particular connections to memory formation after training with feedback.

## Summary

In summary, we found that reward and punishment have distinct effects on resting-state functional connectivity due to training. This suggests that rather than a critical feedback-processing network, feedback may be processed within those systems specifically engaged by the task requirements.

## Supplemental information

**Supplemental figure 1.**
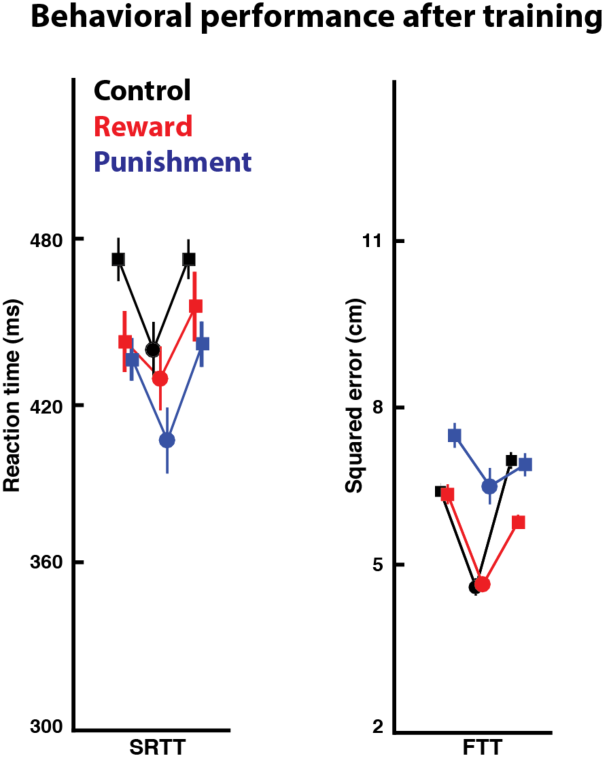
Behavioral performance after training on the SRTT (left) and FTT (right). Participants performed one random sequence block (first square), one fixed-sequence block middle circle), and one random sequence block (second square). During the final three blocks of training on the SRTT, punishment had the best performance regardless of block type (lowest median reaction time). In contrast, during the FTT, punishment performed worst regardless of block type (highest median squared error). A full, quantitative description and statistical analysis of the behavioral results is available in Steel et al. 2018.

## Acknowledgements

AS and CIB are funded by the NIMH internal research program (ZIA-MH002893). CJS holds a Sir Henry Dale Fellowship, funded by the Wellcome Trust and the Royal Society (102584/Z/13/Z). The Wellcome Centre for Integrative Neuroimaging is supported by core funding from the Wellcome Trust (203139/Z/16/Z). The authors would like to thank Matthew Rushworth for his helpful comments on the manuscript.

